# Dissociation of putative open loop circuit from ventral putamen to motor cortical areas in humans I: high-resolution connectomics

**DOI:** 10.1101/2024.08.28.610129

**Authors:** Elizabeth Rizor, Neil M. Dundon, Jingyi Wang, Joanne Stasiak, Taylor Li, Andreea C. Bostan, Regina C. Lapate, Scott T. Grafton

## Abstract

Human movement is partly organized and executed by cortico-basal ganglia-thalamic closed-loop circuits (CLCs), wherein motor cortical areas both send inputs to and receive feedback from the basal ganglia, particularly the dorsal putamen (PUTd). These networks are compromised in Parkinson’s disease (PD) due to neurodegeneration of dopaminergic inputs primarily to PUTd. Yet, fluid movement in PD can sporadically occur, especially when induced by emotionally arousing events. Rabies virus tracing in non-human primates has identified a potential alternative motor pathway, wherein the ventral putamen (PUTv) receives inputs from subcortical limbic areas (such as amygdala nuclei) and sends outputs to motor cortical areas putatively via the nucleus basalis of Meynert (NBM). We hypothesize that this separable open loop circuit (OLC) may exist in humans and explain the preservation of movement after CLC degradation. Here, we provide evidence for the normal human OLC with ultra-high field (7T), multi-echo functional magnetic resonance imaging. We acquired resting-state functional connectivity (FC) scans from 21 healthy adults (avg. age = 29, 12M/9F, all right-handed) and mapped left-hemisphere seed-to-voxel connectivity to assess PUTv FC with putative subcortical nodes and motor cortical areas. We found that putative OLC node (basolateral amygdala, NBM) FC was greater with PUTv (p < 0.05), while CLC subcortical seed (ventrolateral nucleus of thalamus) FC was greater with PUTd (p<0.01). Striatal FC patterns varied across cortical motor areas, with cingulate (p < 0.0001) and supplementary (p < 0.0001) motor areas showing greater FC with PUTv vs. nucleus accumbens. SMA had greater FC with PUTd vs. PUTv (p < 0.0001), while cingulate and primary motor areas showed no significant differences in FC between PUTd and PUTv (p > 0.1). Collectively, these results suggest that PUTv is functionally connected to motor cortical areas and may be integrated into a separable motor OLC with subcortical limbic inputs.

## Introduction

Traditional neuroanatomical understanding of movement control in primates centers on reciprocal projections between motor areas and the basal ganglia, also known as “closed-loop” circuits (CLC) (1). Canonical CLCs associated with movement initiation and inhibition consist of parallel corticostriatal circuits, where motor cortical areas (e.g., cingulate motor area [CMA], supplementary motor area [SMA], primary motor cortex [M1]) input into select striatal regions, whose output projections terminate back in the same motor cortical areas by way of the ventrolateral thalamus (VL). CLCs contain both direct and indirect pathways; both enter the basal ganglia via motor striatum (dorsal putamen [PUTd]), which receives dopaminergic input from the substantia nigra pars compacta (SNpc) (2). The direct CLC routes through the internal segment of the globus pallidus (GPi) and back to the respective motor cortical area (3). The indirect CLC routes back to motor cortical areas via outputs to the external segment of the globus pallidus (GPe) and subthalamic nucleus, which then project to GPi. In both pathways, VL serves as a relay station for GPi outputs to cortical areas. These parallel circuits assist with planning and coordinating movements on finer spatio-temporal scales by integrating sensory and motor information.

It follows that Parkinson’s disease (PD)—a disease characterized at least in its earlier stages by targeted depletion of dopaminergic inputs from SNpc to PUTd—leads to profound and varied movement deficits, such as bradykinesia (slowness of movement), akinesia (absence of movement) rigidity (muscle stiffness) and dyskinesia (involuntary movements) (4). However, both transneuronal-tracer and neurochemical evidence suggest the presence of an alternative circuit through basal ganglia that might be spared by the disease. First, retrograde transneuronal transport of rabies virus in both nonhuman primates has revealed that ventral putamen (PUTv), a subregion of the ventral (limbic) striatum, sends projections to primary motor cortex (M1), but does not receive them in turn (5–8). These findings suggest that a limbic “open-loop” circuit (OLC) might link PUTv to motor areas alongside the CLCs. The primate ventral striatum (PUTv and nucleus accumbens [NAc]) is not known to participate in CLC pathways via projections to GPi (7, 9); instead, it sends outputs primarily to the ventral pallidum and the nucleus basalis of Meynert (NBM), whose cholinergic outputs widely target the cerebral cortex (10–12). The ventral striatum also receives subregion-dependent input from nuclei in the amygdala (PUTv: basolateral nuclei, NAc shell: central, periamygdaloid and basolateral nuclei) (13, 14). As such, we hypothesize that the basolateral amygdala (PUTv input) and NBM (PUTv output) serve as candidate “nodes” of this putative OLC.

Second, PUTv shows spared dopaminergic innervation in primate analogues of PD (15), consistent with observations in humans at the early stage of the disease, where dopamine-agonizing medications seemingly disrupt motor function by way of an “overdose” effect, particularly when sites of spared innervation (PUTv) are the principal task correlates (16). Another phenomenon occasionally seen in those with PD is “Paradoxical Kinesia” (PK) (17), whereby emotionally arousing circumstances such as car accidents (18) or natural disasters (19, 20) can seemingly drive transient normalization of motor function (21).

Together, these findings suggest that the putative OLC might subserve limbic and arousal processes access to the control of movement in certain contexts. Here, we provide ultra-high field (7T) connectomic evidence with multi-echo resting-state functional magnetic resonance imaging (MRI) that a homologous circuit, as seen in primates, might indeed exist in humans. We additionally reveal a connectomic profile that supports the above clinical hypothesis, i.e., a potential circuit in healthy participants that affords communication between limbic/arousal sites and motor areas, specifically via PUTv (the striatal region less affected by PD).

## Results

Twenty one healthy human participants (average age = 29 years, 12M/9F, all right-handed) participated in an ultra-high field (7T), multi-echo resting-state functional connectivity (FC) MRI experiment, contributing a single ∼10-minute scan each. While targeted imaging of the basal ganglia is highly prone to signal loss, multi-echo imaging at 7T has been shown to enhance putamen signal quality; furthermore, 10-minute acquisitions have shown equivalent efficacy when compared to 30-minute, single-echo acquisitions (22, 23). We combined the signal across three echoes and parsed BOLD from non-BOLD components with a data-driven, multi-echo independent component analysis (ME-ICA) approach (Tedana, see Methods). Neural time series were further corrected for physiological noise (recorded respiration and pulse). For each hypothesis-relevant seed region, we calculated the seed average BOLD time series in participant native anatomical space, and quantified seed-to-voxel connectivity using Fisher’s Z transformations of Pearson correlations. To examine evidence for the putative OLC in humans, our analyses had two aims: 1) Assess which putamen subregion(s) (PUTv, PUTd) were most functionally connected to putative OLC nodes (BLA, NBM) and a key CLC subcortical region: ventrolateral thalamus (VL), and 2) Assess whether PUTv possessed functional connectivity to motor cortical areas (SMA, CMA, M1 upper limb area [M1_ul_]) as compared to another limbic striatal subregion (NAc) and PUTd, a known component of the classic motor CLC (**Fig. 1**). We conducted analyses with left hemisphere seeds, as our participant pool was right-handed.

**Fig. 1.**
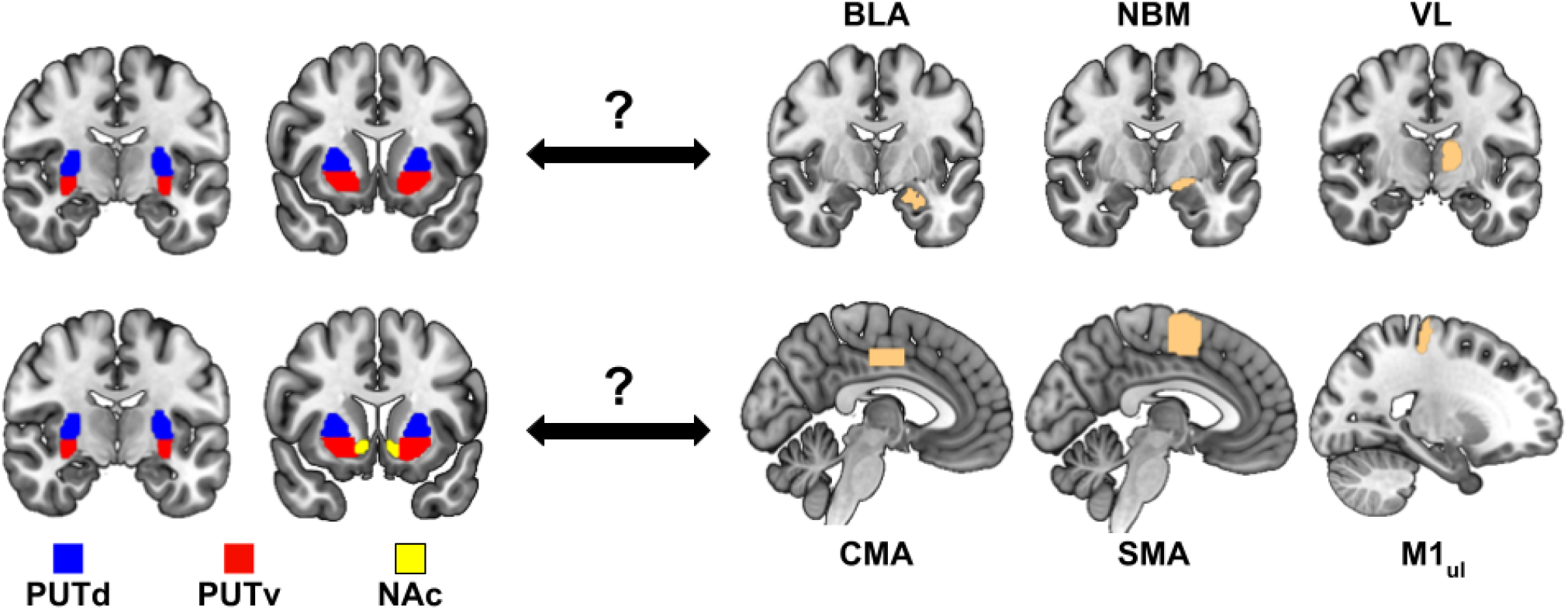
Connectomic Aims. Top row: Connectomic aim 1; Which voxels in the putamen show greatest connectivity to the putative subcortical OLC (BLA, NBM) vs. control CLC (VL) seed regions? Bottom row: Connectomic aim 2; How does PUTv FC with cortical motor areas compare with that of classic motor (PUTd) vs. limbic (NAc) striatal subregions?

### Aim 1: Subcortical Node Seed-Putamen Voxel FC

We first sought connectomic evidence in support of the hypothesized subcortical nodes of the putative OLC, that is, ventral putamen projections to and from sites relevant for limbic and arousal function (Aim 1). To this end, we used seed regions in basolateral amygdala (BLA) and basal forebrain (nucleus basalis of Meynert—NBM).

In order to assess whether putamen subregions (PUTv, PUTd) were significantly more connected with OLC subcortical areas when compared to a CLC control region (VL), we performed two, two-way repeated-measures ANOVAs [(PUTv vs. PUTd) × (OLC vs. CLC)], one for BLA, and one for NBM as OLC nodes. When examining BLA as an OLC node, we identified a significant interaction effect (F = 28.65, p < 0.0001, η^2^ = 0.050); main effects analyses revealed that the putamen subregion did not have a statistically significant effect on mean FC (p = 0.38), but the subcortical node (BLA vs. VL) did (p < 0.0001). When examining pairwise tests in greater detail (**Fig. 2)**, we found that BLA is more connected with PUTv than PUTd (p < 0.05, Cohen’s d = 0.32), whereas VL is more connected with PUTd than PUTv (p < 0.01, d = 0.61)

**Fig 2.**
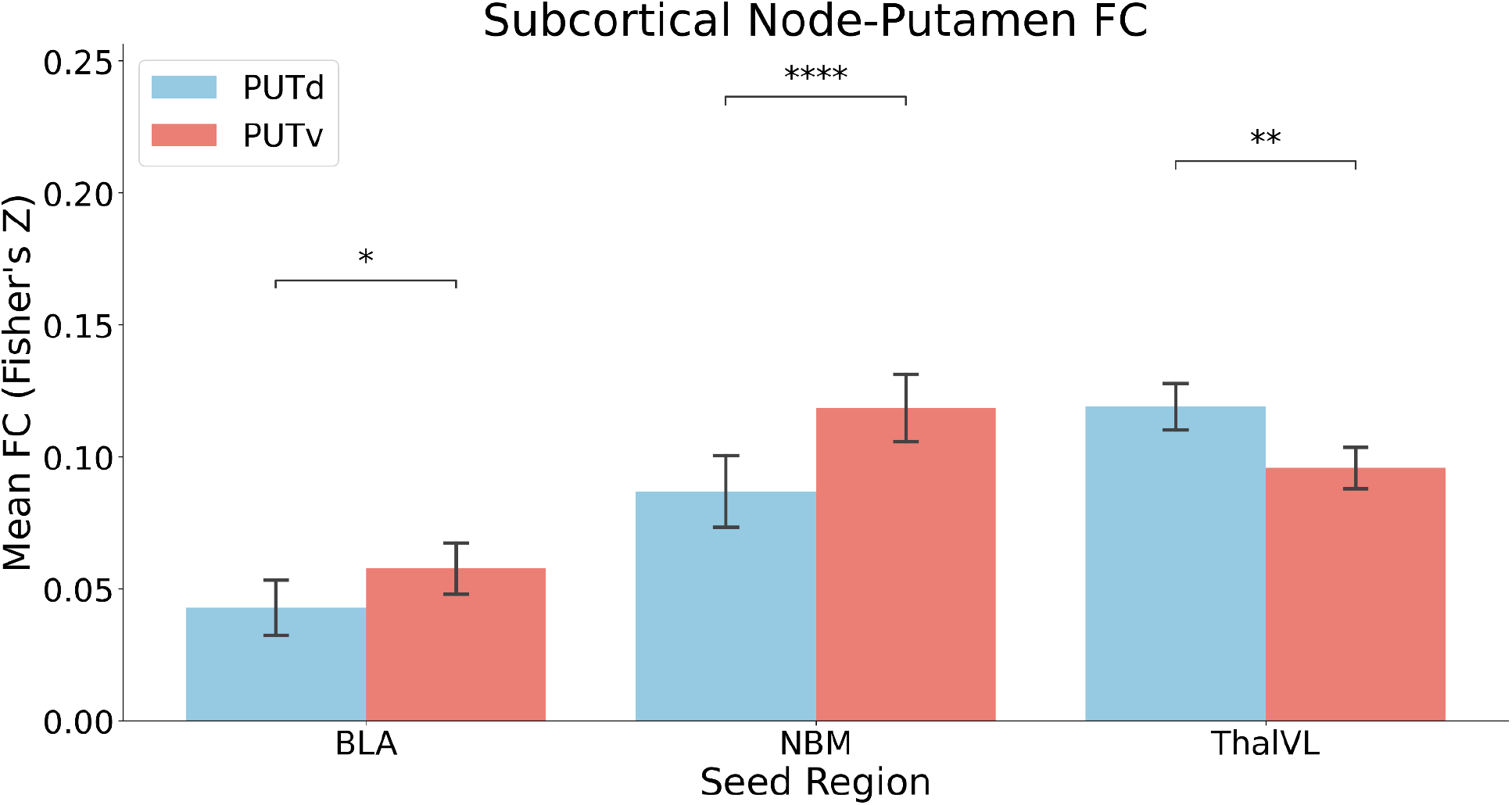
Differences in PUTd vs. PUTv functional connectivity with putative OLC and CLC subcortical node seeds. Legend: * indicates p<0.05, ** indicates p<0.01, **** indicates p<0.0001, ns indicates no significance.

When examining NBM, we again found a significant interaction effect (F = 102.47, p < 0.0001, η^2^ = 0.07); however, no main effects were significant. As we found with BLA, pairwise effects revealed that NBM is more connected with PUTv than PUTd (p < 0.0001, d = 0.52), while, as stated above, VL is more connected with PUTd than PUTv.

**Figure 3** displays how group-averaged striatal FC with subcortical seeds (BLA, NBM, VL) differs across putamen subregions. Connectivity peaks for BLA are found primarily in ventrolateral areas; NBM peaks are also in ventral putamen, but in more rostral areas when compared to BLA. Using a CLC control seed region in VL, we instead see greater PUTd vs. PUTv connectivity in rostral areas and more evenly distributed FC in caudal areas.

**Fig 3.**
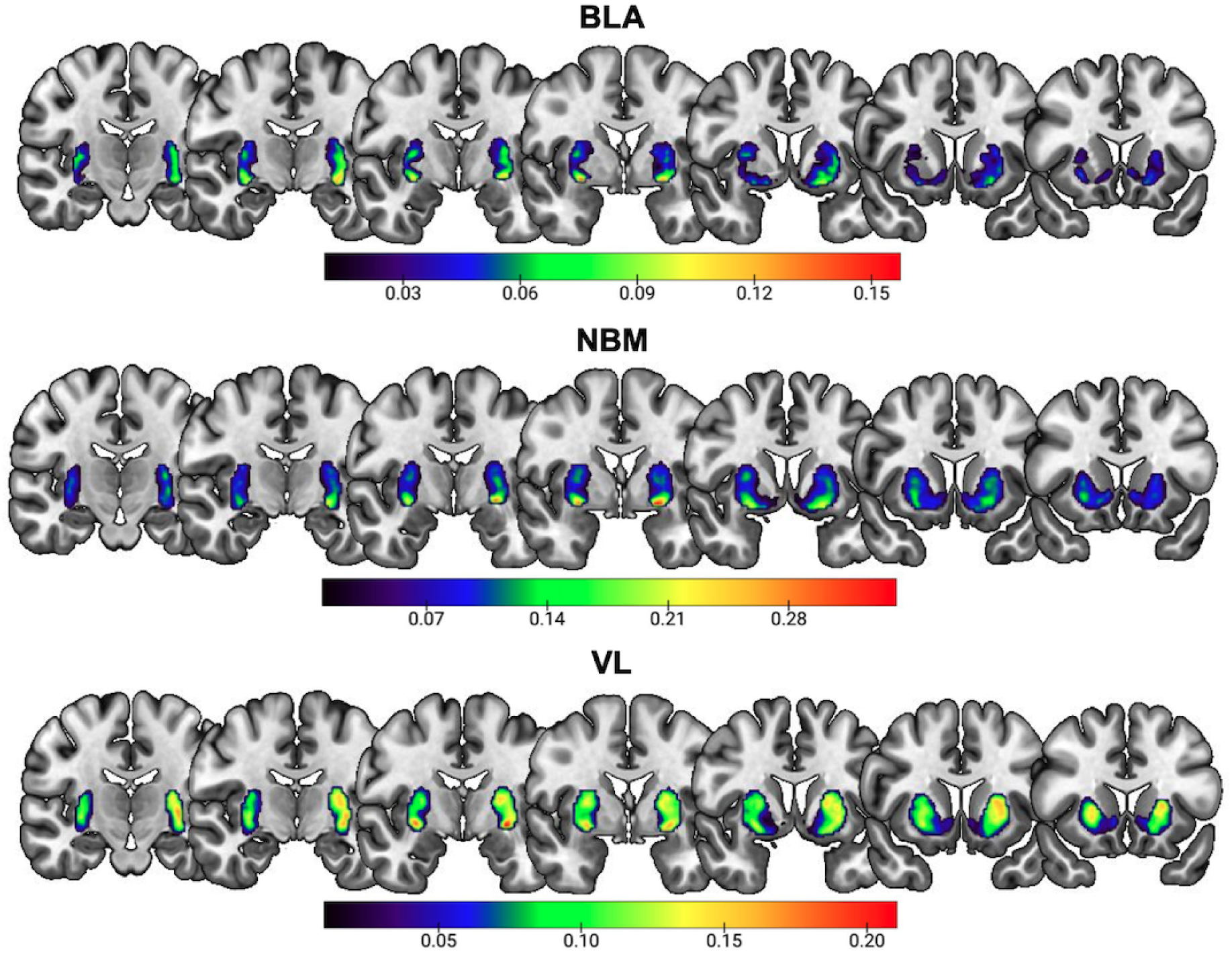
Seed (BLA, NBM, VL) to voxel (PUT + NAc) connectivity maps averaged across all participants (N=21). Color bars indicate the magnitude of Fisher’s Z score ranging from 0.01 (dark purple) to seed-specific relative maximum connectivity value (red). Coronal slices are placed at, from viewer left to right, y=-15, -10, -5, 0, 5, 10, 15.

Overall, we found that PUTd vs. PUTv have distinct FC profiles with hypothesized OLC and CLC subcortical nodes. These findings support the hypothesis that OLC sites relevant for limbic and arousal functions are more connected to the putamen via its ventral division.

### Aim 2: Motor Cortical Seed-Striatal Voxel FC

We next sought connectomic evidence in support of the hypothesized terminals of the putative OLC, that is, projections from PUTv to motor areas. In order to assess whether PUTv was significantly connected with motor cortical areas when compared to motor striatal (PUTd) and limbic striatal (NAc) control regions, we performed two, two-way repeated-measures ANOVAs: [(PUTv vs. PUTd) × (CMA vs. SMA vs. M1_ul_)] and [(PUTv vs. NAc) × (CMA vs. SMA vs. M1_ul_)]. When examining PUTv vs. PUTd, we identified a significant interaction effect (F = 14.78, p < 0.0001, η^2^ = 0.020); main effects analyses revealed that both cortical motor area (p < 0.0001) and putamen subregion (p < 0.001) had statistically significant effects on mean FC. Pairwise tests (**Fig. 4)** revealed that PUTv and PUTd were not differentially connected with CMA or M1_ul_ (p > 0.1); however, PUTd was significantly more connected to SMA than PUTv (p < 0.0001, d = 0.98).

**Fig 4.**
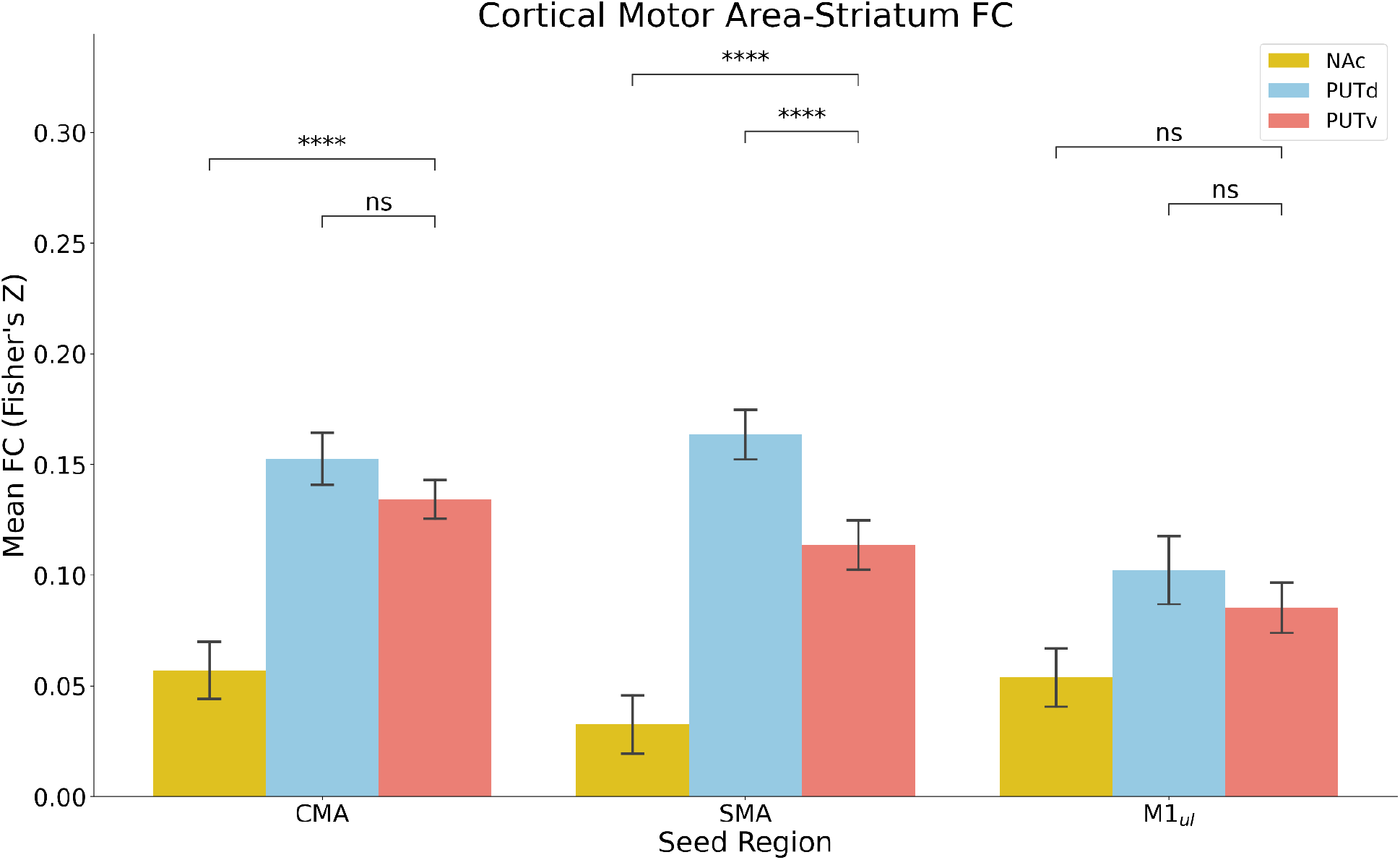
Differences in PUTv vs. motor (PUTd) and limbic (NAc) striatum FC with motor cortical area seeds. Legend: **** indicates p<0.0001, ns indicates no significance.

When examining PUTv vs. NAc, we also identified a significant interaction effect (F = 10.78, p < 0.001, η^2^ = 0.043); main effects analyses also revealed that both motor cortical area (p < 0.01) and striatal subregion (p < 0.0001) had significant effects on mean FC. Pairwise tests (**Fig. 4)** revealed that PUTv trended toward greater FC with M1_ul_ than NAc; however, the test did not pass multiple-comparison correction (p = 0.056, d = 0.27). On the other hand, PUTv was significantly more connected to SMA (p < 0.0001, d = 1.45) and CMA (p < 0.0001, d = 1.52) than NAc. Lastly, to ensure that NAc does not simply have overall weaker connectivity than putamen subregions, especially PUTd, we examined striatal FC with the medial prefrontal cortex (mPFC) with a one-way repeated-measures ANOVA: [mPFC × (NAc vs. PUTv vs. PUTd)]. The mPFC is a key region in reward circuitry with previously observed projections to/connectivity with NAc (8, 24). We identified a significant difference in FC across the 3 striatal subregions (F = 6.26, p < 0.01, η^2^ = 0.11); pairwise tests revealed that PUTd had significantly less FC with mPFC when compared to NAc (p < 0.05, d = 0.78) and PUTv (p < 0.05, d = 0.41). On the other hand, PUTv and NAc were not differentially connected with mPFC (p > 0.25).

Finally, **Figure 5** displays how group-averaged striatal FC with motor cortical seeds (CMA, SMA, M1_ul_) differs across putamen subregions. Both CMA and M1 have similar connectivity profiles, with peaks in midline vPUT and caudal dPUT. In contrast, SMA FC more strongly peaks in dPUT, especially in rostral areas, when compared to CMA and M1.

**Fig 5.**
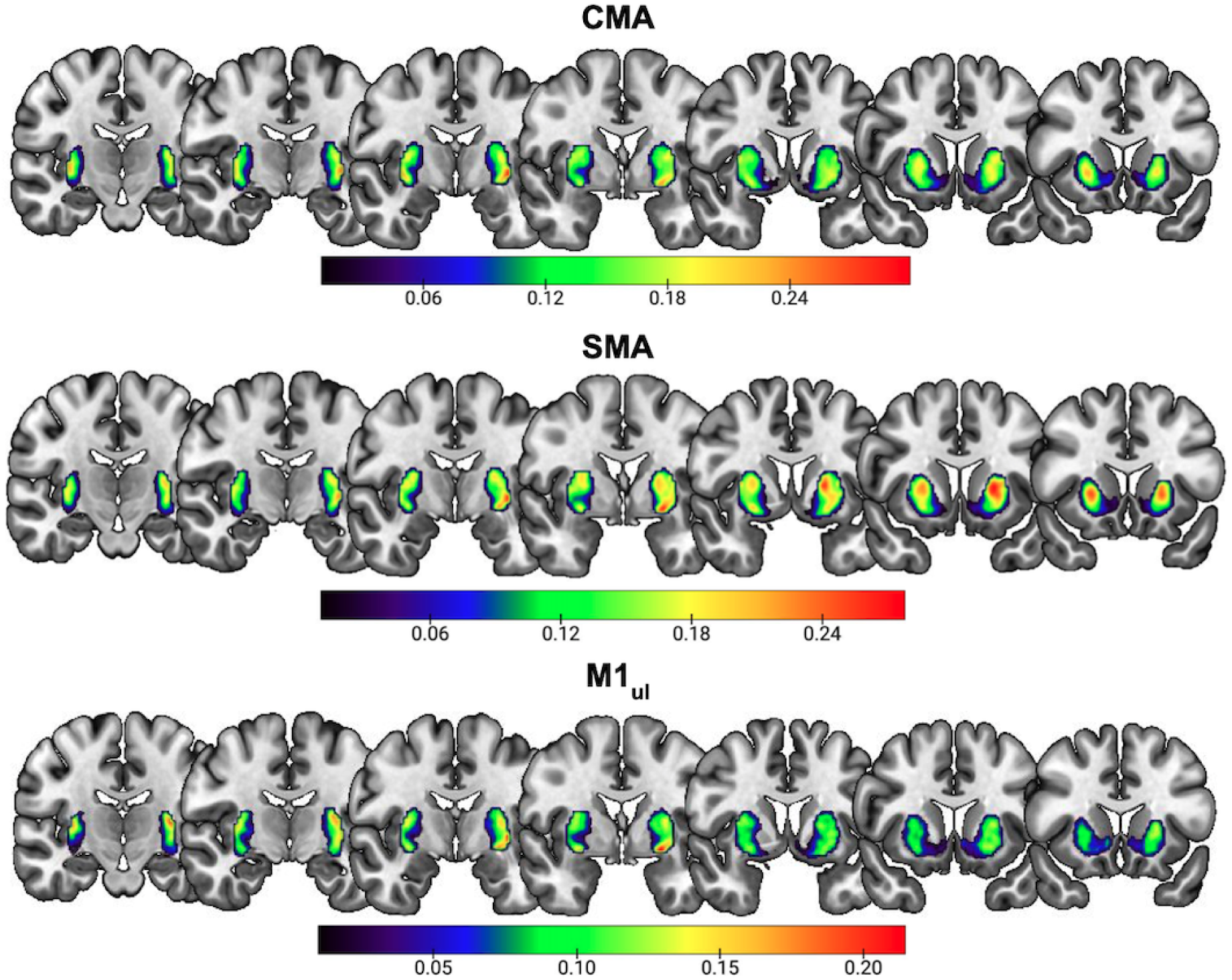
Seed (CMA, SMA, M1_ul_) to voxel (PUT + NAc) connectivity maps averaged across all participants (N=21). Color bars indicate the presence of Fisher’s Z score ranging from 0.01 (dark purple) to seed-specific relative maximum connectivity value (red). Coronal slices are placed at, from viewer left to right, y=-15, -10, -5, 0, 5, 10, 15.

**Table 1.**
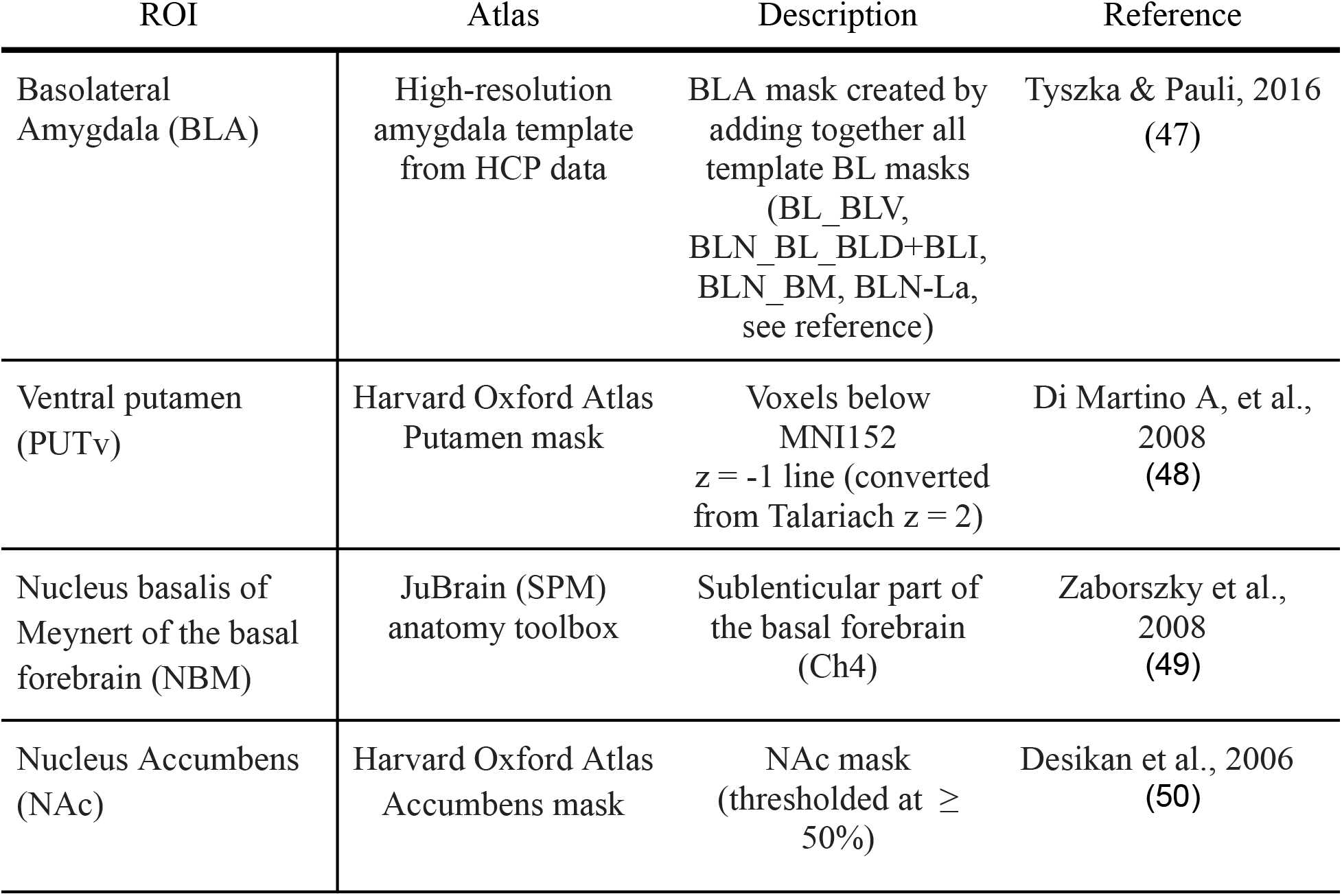
Open Loop Circuit/Ventral striatum ROIs.

**Table 2.** Closed Loop Circuit ROIs.

| ROI | Atlas | Description | Reference |
| --- | --- | --- | --- |
| Dorsal putamen (PUTd) | Harvard Oxford Atlas Putamen mask | Voxels above MNI152 $z = -1$ line (converted from Talariach $z = 2$) | Di Martino A, et al., 2008 (23) |
| Ventrolateral Nucleus of Thalamus (VL) | Automated Anatomical Labeling Atlas 3 | Thal_VL nucleus mask | Rolls et al., 2020 (51) |

**Table 3.** Cortical Area ROIs.

| ROI | Atlas | Description | Reference |
| --- | --- | --- | --- |
| Primary motor cortex, upper limb area ($M1_{ul}$) | Brainnetome Atlas | Motor cortex (M1) upper limb area (thresholded at $\geq 25\%$) | Fan et al., 2016 (52) |
| Cingulate Motor Area (CMA) | N/A | Neurologist (senior author STG) hand-drawn based on MNI coordinates relative to midline and location of SMA | Paus, 2001 (53) |
| Supplementary Motor Area (SMA) | Harvard Oxford Atlas | SMA mask (thresholded at $\geq 50\%$) | Desikan et al., 2006 (50) |
| Medial Prefrontal Cortex (mPFC) | Harvard Oxford Atlas | Medial frontal mask (thresholded at $\geq 50\%$) | Desikan et al., 2006 (50) |

Overall, we found that PUTv has greater FC with motor cortical areas than NAc (the traditionally defined limbic striatum); PUTv FC with CMA and M1_ul_ is comparable to that of PUTd (motor striatum). These findings support the idea that PUTv may also project to motor cortical areas and may be integrated into motor network functioning.

## Discussion

Consistent with transneuronal-tracer evidence in nonhuman primates and clinical observations in PD, our data reveal connectomic evidence supporting the existence of a putative OLC through basal ganglia, primarily relayed by PUTv, to regions of motor cortex. While connectomic data are correlational in nature, and cannot inform the directionality of projections, our data nonetheless reveal plausible communication between key regions of OLC that justify further study.

### Putamen Subcortical Connectivity

We reveal first that sites relevant for limbic and arousal function are primarily associated with the ventral region of putamen. In particular, we saw that putative OLC nodes BLA and NBM were significantly more associated with PUTv when compared to PUTd. When examining average FC patterns across all participants, we see that BLA FC with PUTv spans along the rostrocaudal axis, similarly to patterns previously observed in non-human primates (13). NBM FC, in contrast, is stronger in more rostral regions than that of BLA. Previous work demonstrated that rostral PUTv sends substantial projections to NBM; however, caudal striatum was not examined (10). Conversely, thalamic (CLC) FC was ultimately more associated with PUTd, though hotspots were observed in ventral areas as well. This finding is consistent with non-human primate work showing that VL mainly projects to dorsal and central striatum, while midline thalamic nuclei (not examined here) project more to ventral/limbic areas (25).

Overall, our subcortical connectivity results complement existing observations of a dorsal/ventral striatum gradient in human non-human primates (8, 26–28), providing evidence for amygdala/NBM involvement in a putative OLC. Generally, these findings report that ventral striatum associates with subcortical areas relevant for affective and motivational processes, while dorsal striatum instead associates with subcortical areas relevant for motor, cognitive and executive areas. However, human FC data occasionally deviates from classic NHP anatomical patterns. Notably, Jung et al., 2014 reported strong significant amygdala activity across the putamen; this finding contrasts with NHP tracing studies showing concentrated amygdala connectivity in ventral striatum (29). While we did find greater amygdala FC with PUTv vs.

PUTd, our observed difference was smaller than hypothesized and may reflect these previously observed FC patterns. Future investigation is needed to determine whether this deviation is reflective of true connectivity or is due to fMRI limitations such as signal dropout or bleeding.

### Putamen Cortical Motor Area Connectivity

While our subcortical results generally align with traditional understanding of striatal dorsal/ventral gradients, our motor cortical results paint a more complex, region-dependent picture. As expected, SMA showed stronger FC with PUTd vs. PUTv; we also observed increased group-averaged SMA FC primarily in rostral, dorsomedial areas, as opposed to increased M1_ul_ FC in more caudal, dorso-/ventrolateral areas. This pattern appears consistent with previous reports that SMA projects to areas medial of M1 targets (30, 31). In contrast, PUTv and PUTd were similarly connected with CMA and M1_ul_, providing evidence that PUTv may also be integrated into cortical motor networks, possibly due to OLC pathways. While our observed group-averaged M1_ul_ FC patterns were similar to previous findings, with M1 distal forelimb targets evolving from ventrolateral areas in rostral PUT to dorsolateral areas in caudal PUT, our statistical findings deviate from the expected striatal gradient structure for M1_ul_ and CMA inputs (6, 31). Striatal inputs from cortex are traditionally considered to be organized along a functional axis, where motor inputs target dorsolateral regions, associative inputs target central regions, and limbic inputs target ventromedial regions; thus, PUTv is not known to receive motor cortical inputs (8, 32, 33). Our results also differ from previous fcMRI work at 3T showing dorsal/ventral gradient differences in striatal-M1 FC, with anterior-ventral putamen lacking connectivity to M1 (27, 28). However, previous NHP work indicates that an “open loop” structure may exist where PUTv outputs reach motor cortical areas by way of intermediate regions (5). While fcMRI cannot distinguish connection directionality, our results may reflect this putative OLC output (as opposed to input) connectivity that may be more salient when imaging at 7T.

Our results are possibly region-dependent due to functional differences across cortical motor areas. PUTd’s stronger connectivity with SMA is perhaps consistent with SMA’s involvement with motor planning, especially with regard to sequences of highly learned actions (34). In contrast, CMA (with which we observed comparable associations with PUTd and PUTv) is instead thought to facilitate inner drives, motivational and affective responses (35). Human neuroimaging investigations have identified diverging trajectories for SMA, M1, and CMA in PD, with both decreased SMA-putmen FC and SMA task activation, increased SMA-M1/M1-motor network FC, and lack of significant observed change in CMA (36–38). Though we studied young, healthy controls, our results in context suggest that the putative OLC may rely more on projections to M1 and CMA (balanced FC in dorsal vs. ventral striatum) as PUTd-SMA connection integrity becomes compromised in PD. Our results also suggest that the classic dorsal/ventral gradient may not be a complete description of human striatum-cortex connectivity.

### Motor Cortical Connectivity Profiles within the Ventral Striatum

In order to assess whether FC with motor cortical areas differed between ventral striatal regions, we compared PUTv FC with NAc, another limbic/ventral striatal area. We observed that PUTv was significantly more connected to motor cortical areas than NAc, though the difference in M1_ul_ FC did not pass multiple-comparison correction. As expected, we also observed greater NAc FC with mPFC when compared to PUTd, casting doubt that the identified decreased NAc-motor cortical area FC is simply due to differences in signal strength across the striatum. However, previous NHP work has identified output projections from NAc to M1, as well as increased compensatory NAc-M1 FC after spinal cord injury (6, 39). Recent work in rodents has also proposed the existence of open-loop architecture involving NAc-driven nigrothalamic pathways (40). Therefore, it is possible that NAc-M1 connectivity may be difficult to distinguish from PUTv-M1 connectivity with fMRI. Future investigation is needed to determine whether this proposed nigrothalamic pathway exists in primates, and if so, whether NAc vs. PUTv-driven pathways are anatomically and functionally distinct.

### Limitations and Future Directions

Resting-state functional connectivity imaging has notable limitations, especially when compared to anatomic tracing studies. FC measures cannot distinguish input vs. output projections, nor ascertain how many neurons may mediate connectivity between two regions. Therefore, comparisons to anatomical tracing data must be made cautiously. Additionally, though ultra-high field, multi-echo imaging is known to improve image quality of basal ganglia nuclei, these areas are still susceptible to signal dropout and enhanced noise, potentially affecting interpretation of results. Network topography is known to vary across individuals; therefore, average FC maps may obscure patterns and artificially inflate/deflate FC values. Future work mapping these circuits on an individual basis is warranted.

Our work complements a growing body of work aimed at profiling the connectome of PD. Though we only studied young individuals with no disease diagnosis here, our results serve as a baseline for future mapping of connectivity profile changes in PD. First, it can be determined if corticostriatal connectivity is indeed disrupted between nodes of the CLC affected by Parkinson’s disease, such as PUTd and SMA. Second, it can be inferred whether those with PD exhibit preserved sub-corticostriatal (NBM, BLA with PUTv) and corticostriatal (PUTv/M1_ul_/CMA with PUTv) functional connectivity between putative OLC nodes.

## Conclusion

This study in young, healthy participants reveals high-resolution connectomic evidence that supports a putative OLC through basal ganglia to the motor cortex as one potential substrate. This circuit may be preserved in PD, a disease that affects millions of people worldwide with no known cure. The OLC may serve as a potential target for amplifying movement in PD.

## Glossary

BLA: basolateral amygdala
CLC: closed-loop circuit
CMA: cingulate motor area
fMRI: functional magnetic resonance imaging
GPi: globus pallidus internal segment
M1_ul_: primary motor cortex, upper-limb region
mPFC: medial prefrontal cortex
NBM: nucleus basalis of Meynert of the basal forebrain
OLC: open-loop circuit (putative)
PD: Parkinson’s disease
PK: paradoxical kinesia
PUTd: dorsal (sensorimotor) putamen
PUTv: ventral (limbic) putamen
SMA: supplementary motor area
SNpc: substantia nigra pars compacta
VL: ventrolateral nucleus of the thalamus

## Acknowledgements

We would like to thank Arthur Toga, Danny Wang, Kay Jan, Ioannis Pappas, and Katherin Martin at the USC Mark and Mary Stevens Neuroimaging and Informatics Institute for scan acquisition development and processing assistance. We would also like to thank Logan Dowdle and Dan Handwerker (Tedana team) for acquisition development and Tedana preprocessing guidance. Finally, we want to thank Evan Layher for pilot testing. This research was funded in whole by Aligning Science Across Parkinson’s ASAP-020-519 through the Michael J. Fox Foundation for Parkinson’s Research (MJFF).

## Open Access Policy

All data used in the above analyses are freely available as part of the SKIP (SoCal Kinesia and Incentivization for Parkinson’s Disease) dataset through OpenNeuro.org and available under CC0 public copyright. For further information see www.socalkinesia.org. For the purpose of open access, the author has applied a CC0 public copyright license to all Author Accepted Manuscripts arising from this submission. To access all open source data and code used for this manuscript, please refer to the Key Resource Table.

## Key Resource Table

| RESOURCE TYPE | RESOURCE NAME | SOURCE | IDENTIFIER | NEW/ REUSE | ADDITIONAL INFORMATION |
| --- | --- | --- | --- | --- | --- |
| **Dataset** | SKIP highresFC MRI dataset | OpenNeuro | https://doi.org/10.18112/openneuro.ds005264.v1.0.0 | new | Raw MRI data |
| **Dataset** | FC striatum maps | Zenodo | https://doi.org/10.5281/zenodo.13381546 | new | Group-averaged FC maps for Figs 3,5 |
| **Dataset** | Tabular FC values | Zenodo | https://doi.org/10.5281/zenodo.13381546 | new | Used for stats and Figs 2, 4 |
| **Protocol** | MRI protocol | [protocols.io](https://doi.org/10.17504/protocols.io.dm6gpbj1lzp/v1) | [dx.doi.org/10.17504/protocols.io.dm6gpbj1lzp/v1](https://doi.org/10.17504/protocols.io.dm6gpbj1lzp/v1) | new |  |
| **Software/code** | Preprocessing and analysis scripts | Github/Zenodo | https://doi.org/10.5281/zenodo.10637692 | new |  |
| **Software/code** | Mask repository | Github/Zenodo | https://doi.org/10.5281/zenodo.10637692 | new | ROI masks for FC analysis |
| **Software/code** | FSL 6.0.7.12 | http://www.fmrib.ox.ac.uk/fsl/ | RRID:SCR_002823 | reuse |  |
| **Software/code** | MATLAB R2022a | http://www.mathworks.com/products/matlab/ | RRID:SCR_001622 | reuse |  |
| **Software/code** | TE Dependent ANALysis (Tedana) 24.0.1 | https://tedana.readthedocs.io/en/stable/# |  | reuse |  |
| **Software/code** | Advanced Normalization Tools (ANTs) 2.3.4 | https://github.com/ANTsX/ANTs | RRID:SCR_004757 | reuse |  |
| **Software/code** | PhysIO toolbox 6.0.1 | https://www.nitrc.org/projects/physio/ | RRID:SCR_003430 | reuse |  |
| **Software/code** | Python 3.11.5 | https://www.python.org/downloads/release/python-360/ | RRID:SCR_008394 | reuse |  |
| **Other** | 7T MRI Scanner | Siemens | MAGNETOM Terra | reuse |  |
| **Other** | Head coil | Nova Medical, Inc. | 8-channel pTx system and 8Tx/32Rx head coil | reuse |  |

## Methods

### Participants

A total of 21 young, healthy participants (12M/9F, mean age = 29 years; range = 21-36, all right-handed) completed all study recruitment, pre-screening, and protocol procedures. Study inclusion criteria required that participants be between the ages of 18 – 40 and competent with English enough to understand instructions. Exclusionary criteria were contraindications to MRI (non-removable metal, permanent mouth retainers/braces, incompatible medical devices, vertigo, dizziness, hearing loss/tinnitus, claustrophobia), pregnancy, or clinically significant cardiac, neurological, pulmonary, or psychiatric diseases. Participants were recruited via word of mouth and digital flyers sent to the University of California, Santa Barbara and University of Southern California communities. To assess eligibility, participants completed online screening forms that assessed demographics, health history, and MRI safety eligibility. All participants provided written informed consent for study procedures approved by the Institutional Review Boards at University of California, Santa Barbara and the University of Southern California and were paid

$20/hour for the full MRI session. Data were acquired at the Center for Image Acquisition at the University of Southern California Mark and Mary Stevens Neuroimaging and Informatics Institute.

### Study Protocol

On the day of their study session, participants completed the Edinburgh Handedness Inventory (41) and a comprehensive questionnaire assessing demographics, medication use, and health history. Participants then completed a 30-minute MRI protocol with at the University of Southern California’s Center for Image Acquisition, where they were scanned with a Siemens 7T Terra scanner and a research-only 8-channel transmit 32-channel receive Nova 8Tx/32Rx head coil (Nova Medical Inc.)

First, high-resolution T1-weighted magnetization prepared rapid gradient echo (MPRAGE) anatomical scans were acquired (TR = 4300 ms, TE = 2.27 ms, FOV = 240 mm, T1 = 1000 ms, flip angle = 4°, with 0.75 mm^3^ voxel size). Following the anatomical scan, modified CMRR multi echo resting-state echo planar imaging (EPI) sequences were acquired (Total time = 10:52 mins, TR = 2100ms, flip angle = 70°, 1.5mm thick axial slices, 2×2 mm in plane, 300 volumes, 200mm FOV, iPat GRAPPA factor = 3, MB factor = 4, phase encoding = posterior to anterior), where participants were instructed to keep their eyes open. The echo times and slice number were slightly modified for the first 8 participants (TE1=15.20ms, TE2=34.23ms, TE3=53.26ms, 84 slices) vs. the remaining 13 (TE1=15.20ms, TE2=33.87ms, TE3=52.54ms, 100 slices). Two spin-echo EPI sequences for field map generation were also acquired: one with anterior to posterior (AP) and another with posterior to anterior (PA) phase encoding to use for distortion correction (TR = 3000ms, TE = 60ms, flip angle = 70°, 1.5mm thick axial slices, 2×2mm in plane, 192mm FOV, iPat GRAPPA factor = 2, MB factor = 5).

During scanning, participant respiration and pulse measurements were acquired with an in-scanner Siemens respiration belt and pulse sensor for later denoising. Testing of one participant (not included in the study sample) found that the modified CMRR multi echo EPI sequence utilized for this study produced more optimal temporal signal-to-noise ratio (tSNR) values when compared to other options; further details can be found in the Supplementary Information (p.1).

### Magnetic Resonance Imaging Preprocessing

We conducted MRI preprocessing with a custom pipeline featuring Advanced Normalization Tools (42), FSL (43), TE Dependent ANAlysis version 24.0.1 (44), and custom scripts written in MATLAB (45). Our preprocessing pipeline was inspired by Lynch et al., 2020 (22). First, T1-weighted anatomical data were skull-stripped with antsBrainExtraction.sh and resampled to functional image resolution (2mm voxels). We then used FSL Topup with AP and PA spin echo sequences to obtain an undistorted magnitude image and fieldmap for distortion correction. Next, we averaged the EPI data single band reference images at each timepoint across 3 echoes; this reference image was then brain extracted with FSL BET. EPI data were motion-corrected with FSL MCFLIRT (transforms for echo 1 were applied to echoes 2 and 3) and slice time corrected with FSL slicetimer. We used fsl_motion_outliers to obtain framewise displacement measurements for EPI echo 1 data; 15/21 participants had an average FD of <0.1mm, while 6/21 had an average FD <0.2mm. We used FSL’s epi_reg with the single-band reference brain to obtain fieldmap distortion correction transforms and co-registration transforms to T1-MPRAGE data. These transforms were then applied to EPI data to place all three echoes into participant-specific anatomical space. Due to fieldmap scan acquisition error, the distortion correction transforms were not applied to the first 12/21 participants.

Next, we denoised resting-state EPI data with Tedana. Briefly, Tedana optimally combines the three echo images, reduces data dimensionality with principal component analysis (PCA), and removes TE-independent (non-BOLD) components from the data with independent component analysis (ICA). Tedana pipeline details can be found in the Supplementary Information (p.4). Because subcortical regions are highly susceptible to physiological noise, we then used fsl_regfilt to remove 18 RETROICOR (respiration and cardiac pulse) regressors generated by the PhysIO Toolbox (46). Finally, we smoothed EPI data with a 2.5mm FWHM gaussian kernel; smoothing was minimized due to the subregional focus of this study.

### Region-of-interest Masks

ROI seed masks were deliberately selected from various appropriate atlases for our analyses (Tables 1-3).

### Circuit Connectivity Extraction

For each participant, we used FSL FLIRT to transform all ROI seed masks from MNI152 space to participant-specific anatomical space. Then, for each ROI in participant space, we used custom MATLAB scripts to calculate the ROI’s mean functional data time series, and subsequently calculate a map of Fisher’s Z scores (from Pearson’s r correlation coefficients) between this seed ROI time series and all other voxels in the brain. Fisher’s Z FC values between pairs of ROIs (ROI_1 -ROI_2) were calculated by masking the seed-to-voxel map of ROI_1 with the mask of ROI_2 and averaging voxel-wise Fisher’s Z values within ROI_2’s mask.

Statistical analyses were performed on distributions consisting of FC values calculated in each participant’s individual anatomical space. For group-level visualization (Figures 3, 5), individual seed-to-voxel functional connectivity maps were non-linearly transformed to MNI152 space with antsRegistrationSyN.sh and averaged across all participants.

### Statistical analysis

All statistical analyses were performed with Python (3.11.5). To assess evidence for Connectomic Aims 1 and 2, we conducted two-way repeated measures ANOVAs (pingouin 0.5.4). Effect sizes for ANOVAs were reported using partial eta squared (η^2^; .01-.059 = small effect; .06-.139 = medium effect; ≥.14 = large effect). For Aim 1, we examined the effect of putamen subarea (PUTd, PUTv) on FC with putative OLC (BLA, NBM) and CLC (ThalVL) subcortical seeds. For Aim 2, we examined the effect of putamen subarea (PUTv, PUTd) and limbic striatal subarea (PUTv, NAc) on FC with motor cortical areas (CMA, SMA, M1_ul_).

Post-hoc paired t-tests with Bonferroni correction were also performed to assess FC differences between individual regions. All pairwise test p-values were reported as Bonferroni-adjusted values (i.e., p = 0.05 * 2 regions [BLA/NBM, VL] for Aim 1 and p = 0.05*3 regions [CMA, SMA, M1_ul_] for Aim 2). Lastly, as a control for Aim 2, we examined the effect of striatal subregion (NAc, PUTv, PUTd) on FC with mPFC by performing a one-way repeated measures ANOVA; pairwise p-values for this analysis were also reported as Bonferroni-adjusted values (p = 0.05 * 3 regions [NAc, PUTv, PUTd]). Effect sizes for post-doc paired t-tests were reported using Cohen’s d (d; 0-0.49 = small effect, 0.50-0.79 = medium effect, > 0.80 = large effect).

