## Supplementary Information for "Dissociation of putative open loop circuit from ventral putamen to motor cortical areas in humans I: high-resolution connectomics"

### Supplementary Methods

#### Modified CMRR Scan tSNR

Testing of one participant (not in study sample) found that the modified CMRR multi echo resting-state EPI sequence used for this study had superior tSNR for our regions-of-interest, especially in the amygdala, when compared to two other scan options: 1) the same sequence but with 2mm isotropic voxels, and 2) this sequence modified to have 2mm isotropic voxels and a lower multiband factor (MB=2, TR=3700). The lower-MB sequence was tested due to concerns that an iPat GRAPPA factor of 3 with an MB factor of 4 would induce too much acceleration noise; however, lowering these values drastically increases TR for an acquisition with 1.5x2x2mm voxel size. All three sequences underwent the same preprocessing pipeline, with FSL motion and slice timing correction, followed by denoising with Tedana. tSNR was calculated with fslmaths by dividing the EPI data mean (across time) by the standard deviation. ROI masks were then transformed (to the tSNR maps in participant native anatomic space. Regional mean tSNR was calculated by averaging across all tSNR map voxel values within an ROI mask.

This testing affirmed that raising the acceleration factor to achieve thinner slices at a reasonable TR generally produced greater tSNR than the alternative option. Bar plots below illustrate mean (error bars = SD) tSNR for motor and putative open loop circuit regions.

Basolateral Amygdala

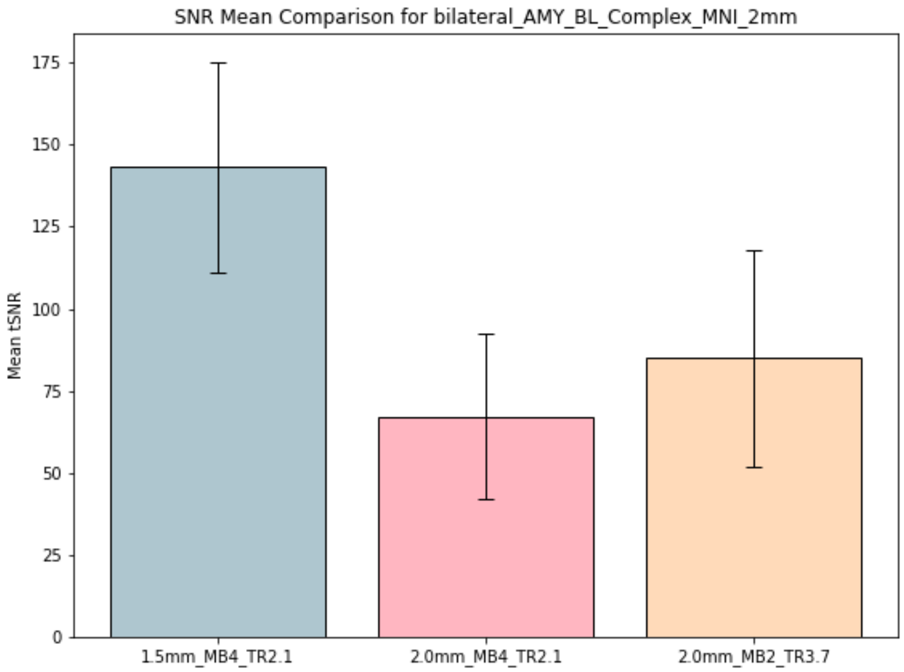

Nucleus basalis of Meynert

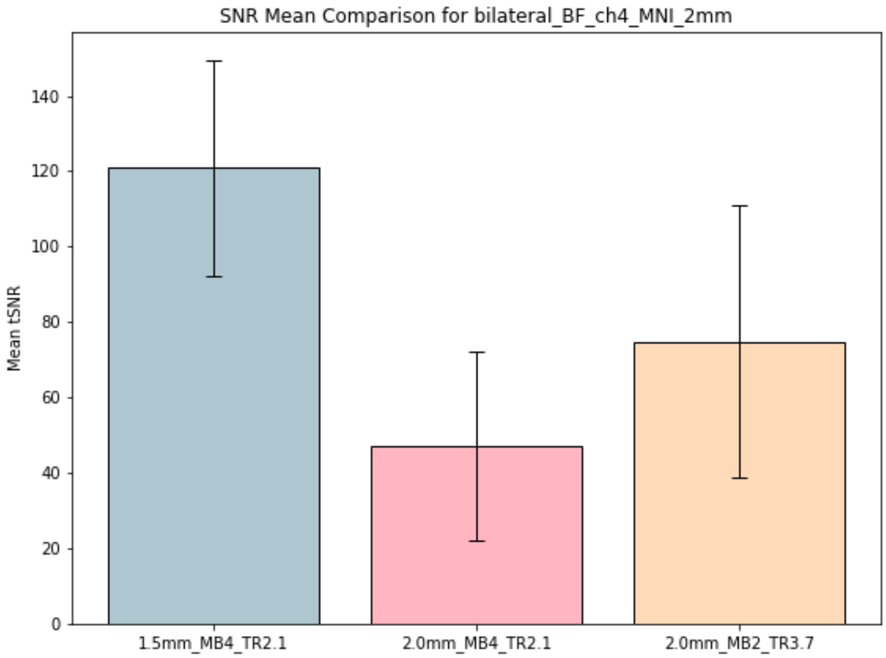


Ventral putamen

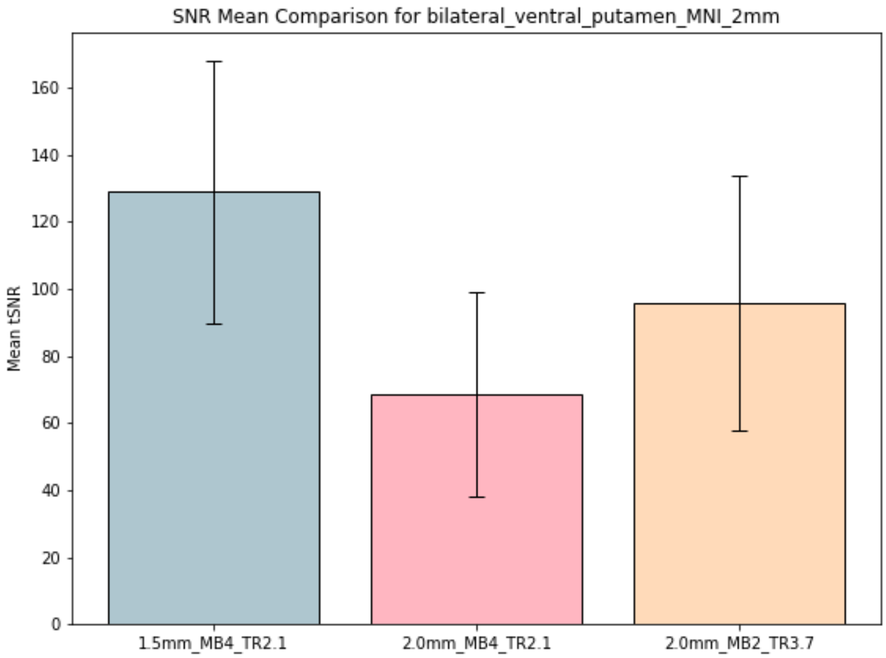

Dorsal putamen

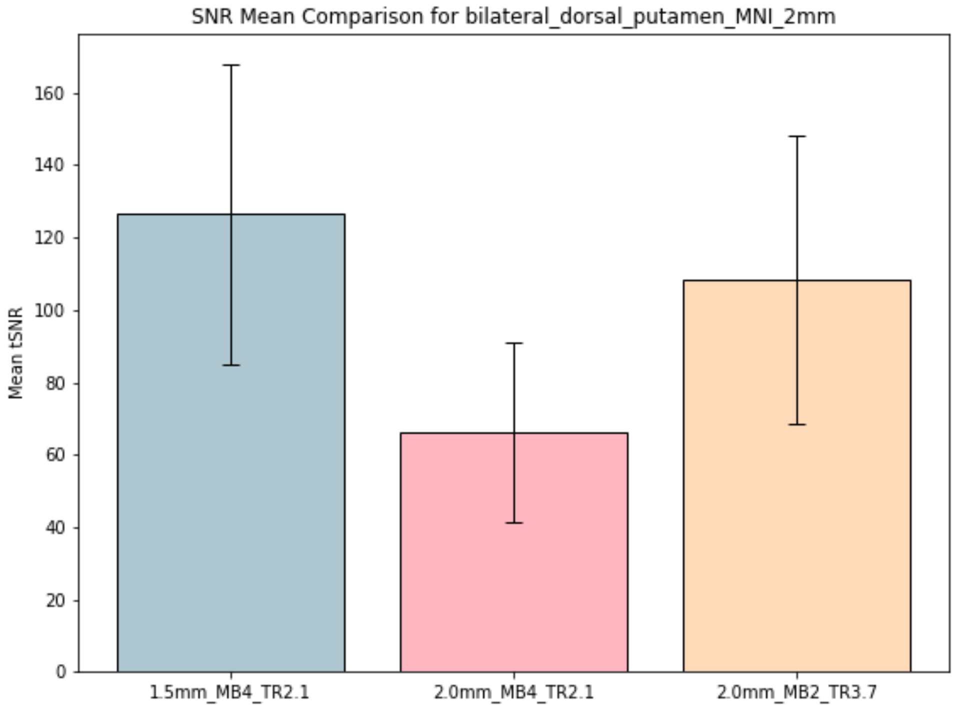


Motor cortex (M1 upper limb)

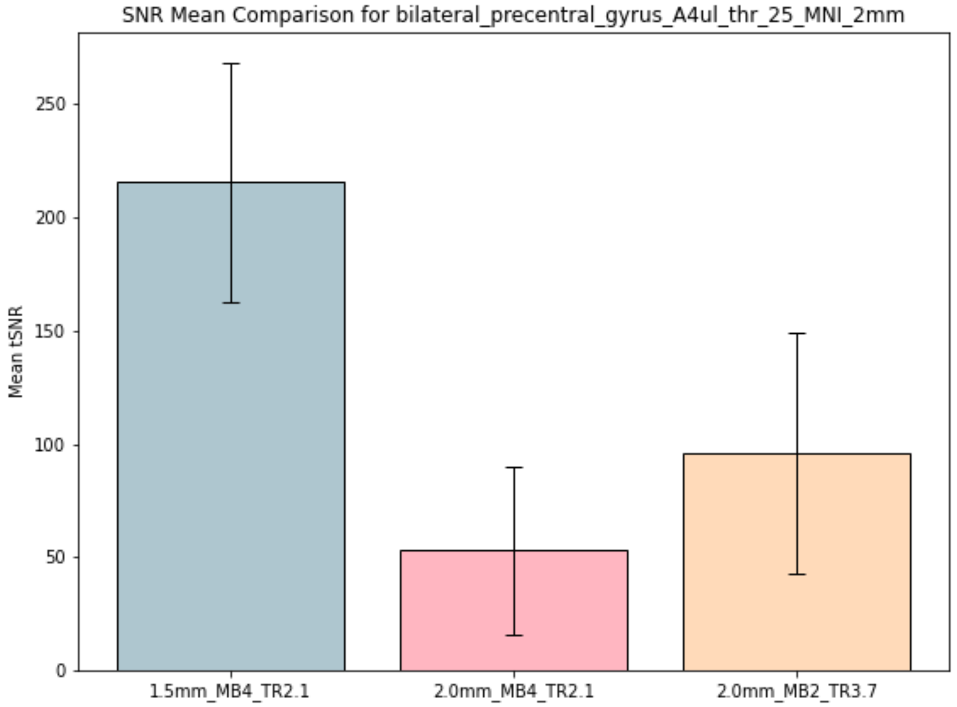

Cingulate motor area (CMA)

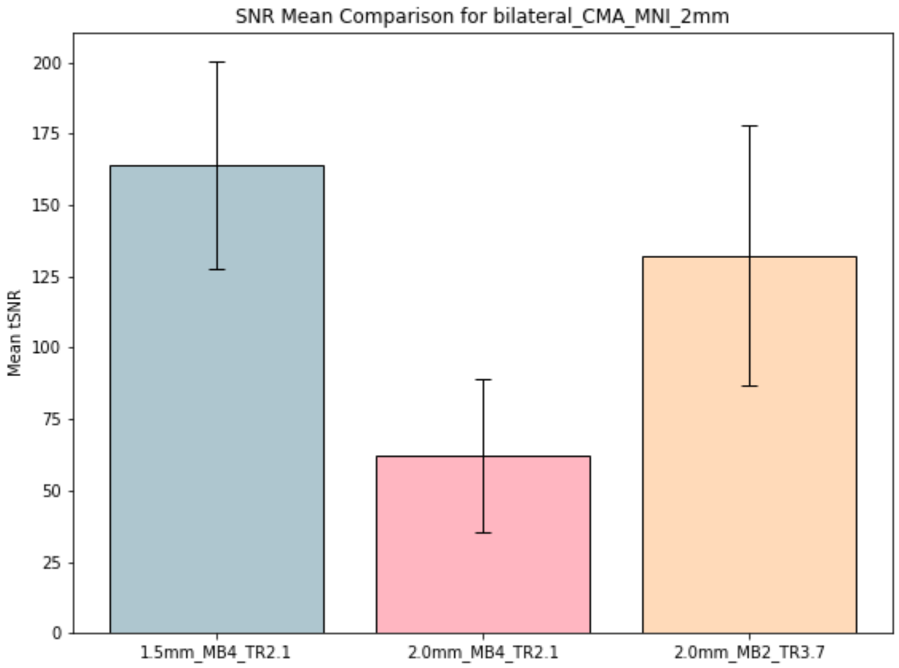


Supplementary Motor Area (SMA)

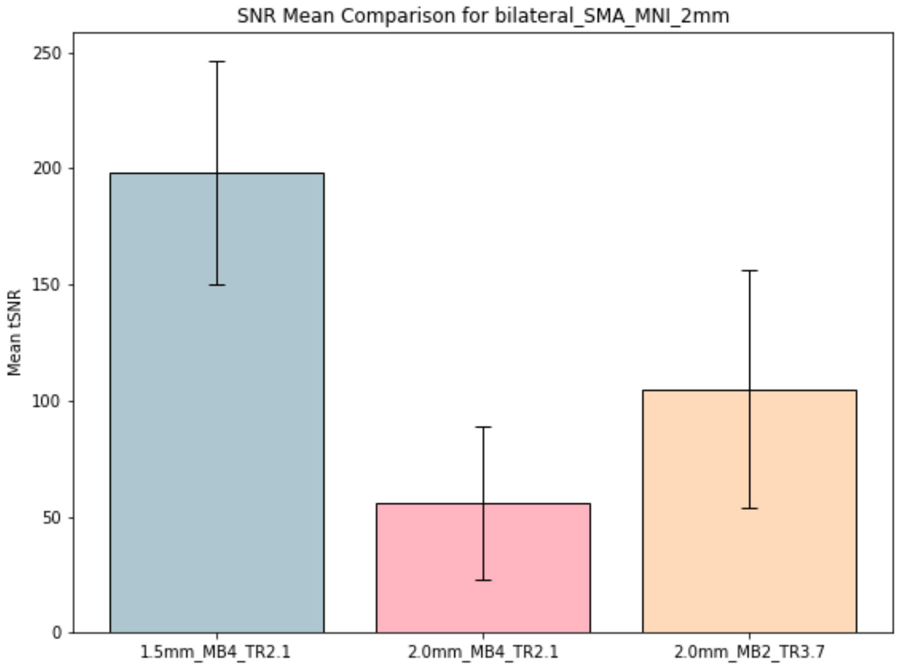

**Tedana methods output**

The ICA decision tree used for this study is the tedana_orig tree (tedana community et al. 2024), which is very similar to the criteria of the MEICA v2.5 decision tree (Kundu et al. 2013). For a description of the decision tree steps, with the rationale for each step, see (Olafsson et al. 2015). TE-dependence analysis was performed on input data using the tedana workflow (DuPre et al. 2021). A user-defined mask was applied to the data. An adaptive mask was then generated using the dropout method(s), in which each voxel's value reflects the number of echoes with 'good' data. An adaptive mask was then generated using the dropout method(s), in which each voxel's value reflects the number of echoes with 'good' data. A two-stage masking procedure was applied, in which a liberal mask (including voxels with good data in at least the first echo) was used for optimal combination, T2*/S0 estimation, and denoising, while a more conservative mask (restricted to voxels with good data in at least the first three echoes) was used for the component classification procedure. A monoexponential model was fit to the data at each voxel using nonlinear model fitting in order to estimate T2* and S0 maps, using T2*/S0 estimates from a log-linear fit as initial values. For each voxel, the value from the adaptive mask was used to determine which echoes would be used to estimate T2* and S0. In cases of model fit failure, T2*/S0 estimates from the log-linear fit were retained instead. Multi-echo data were then optimally combined using the T2* combination method (Posse et al. 1999). Global signal regression was applied to the multi-echo and optimally combined datasets. Principal component analysis in which the number of components is pre-defined was applied to the optimally combined data for dimensionality reduction. The following metrics were calculated: kappa, rho, countnoise, countsigFT2, countsigFS0, dice_FT2, dice_FS0, signal-noise_t, variance explained, normalized variance explained, d_table_score. Kappa (kappa) and Rho (rho) were calculated as measures of TE-dependence and TE-independence, respectively. A t-test was performed between the distributions of T2*-model F-statistics associated with clusters (i.e., signal) and non-cluster voxels (i.e., noise) to generate a t-statistic (metric signal-noise_z) and p-value (metric signal-noise_p) measuring relative association of the component to signal over noise. The number of significant voxels not from clusters was calculated for each component. Independent component analysis was then used to decompose the dimensionally reduced dataset. The following metrics were calculated: countnoise, countsigFS0, countsigFT2, d_table_score, dice_FS0, dice_FT2, kappa, normalized variance explained, rho, signal-noise_t, variance explained. Kappa (kappa) and Rho (rho) were calculated as measures of TE-dependence and TE-independence, respectively. A t-test was performed between the distributions of T2*-model F-statistics associated with clusters (i.e., signal) and non-cluster voxels (i.e., noise) to generate a t-statistic (metric signal-noise_z) and p-value (metric signal-noise_p) measuring relative association of the component to signal over noise. The number of significant voxels not from clusters was calculated for each component.

Next, component selection was performed to identify BOLD (TE-dependent) and non-BOLD (TE-independent) components using a decision tree.

This workflow used numpy (Van Der Walt et al. 2011), scipy (Virtanen et al. 2020), pandas (McKinney et al. 2010, pandas development team et al. 2020), scikit-learn (Pedregosa et al. 2011), nilearn, bokeh (Team et al. 2018), matplotlib (Hunter et al. 2007), and nibabel (Brett et al. 2019). This workflow also used the Dice similarity index (Dice et al. 1945, Sorensen et al. 1948).

Tedana References:


Brett, M., Markiewicz, C. J., Hanke, M., Côté, M.-A., Cipollini, B., McCarthy, P., \u2026 freec84 (2019 , May). nipy/nibabel: 2.4.1.

Dice, L. R. (1945). Measures of the amount of ecologic association between species. Ecology, 26(3), 297\u2013302. URL: https://doi.org/10.2307/1932409, doi:10.2307/1932409

DuPre, E., Salo, T., Ahmed, Z., Bandettini, P. A., Bottenhorn, K. L., Caballero-Gaudes, C., \u2026 others. (2021). Te-dependent analysis of multi-echo fmri with* tedana. Journal of Open Source Software, 6(66), 3669. URL: https://doi.org/10.21105/joss.03669, doi:10.21105/joss.03669

Hunter, J. D. (2007). Matplotlib: a 2d graphics environment. Computing in Science & Engineering, 9(3), 90\u201395. doi:10.1109/MCSE.2007.55

Kundu, P., Brenowitz, N. D., Voon, V., Worbe, Y., Vértes, P. E., Inati, S. J., \u2026 Bullmore, E. T. (2013). Integrated strategy for improving functional connectivity mapping using multiecho fmri. Proceedings of the National Academy of Sciences, 110(40), 16187\u201316192. URL: https://doi.org/10.1073/pnas.1301725110, doi:10.1073/pnas.1301725110

McKinney, W., & others. (2010). Data structures for statistical computing in python. Proceedings of the 9th Python in Science Conference (pp. 51\u201356). URL: https://doi.org/10.25080/Majora-92bf1922-00a, doi:10.25080/Majora-92bf1922-00a

Olafsson, V., Kundu, P., Wong, E. C., Bandettini, P. A., & Liu, T. T. (2015). Enhanced identification of bold-like components with multi-echo simultaneous multi-slice (mesms) fmri and multi-echo ica. Neuroimage, 112, 43\u201351. URL: https://doi.org/10.1016/j.neuroimage.2015.02.052, doi:10.1016/j.neuroimage.2015.02.052

pandas development team, T. (2020 , February). pandas-dev/pandas: Pandas.

Pedregosa, F., Varoquaux, G., Gramfort, A., Michel, V., Thirion, B., Grisel, O., \u2026 others. (2011). Scikit-learn: machine learning in python. the Journal of machine Learning research, 12, 2825\u20132830. URL: http://jmlr.org/papers/v12/pedregosa11a.html

Posse, S., Wiese, S., Gembris, D., Mathiak, K., Kessler, C., Grosse-Ruyken, M.-L., \u2026 Kiselev, V. G. (1999). Enhancement of bold-contrast sensitivity by single-shot multi-echo functional mr imaging. Magnetic Resonance in Medicine: An Official Journal of the International Society for Magnetic Resonance in Medicine, 42(1), 87\u201397. URL: https://doi.org/10.1002/(SICI)1522-2594(199907)42:1<87::AID-MRM13>3.0.CO;2-O, doi:10.1002/(SICI)1522-2594(199907)42:1<87::AID-MRM13>3.0.CO;2-O

Sorensen, T. A. (1948). A method of establishing groups of equal amplitude in plant sociology based on similarity of species content and its application to analyses of the vegetation on danish commons. Biol. Skar., 5, 1\u201334.

Team, B. D. (2018). Bokeh: Python library for interactive visualization. URL: https://bokeh.pydata.org/en/latest/

tedana community. (2024). Component selection decision trees in tedana. figshare. doi:10.6084/m9.figshare.25251433.v2

Van Der Walt, S., Colbert, S. C., & Varoquaux, G. (2011). The numpy array: a structure for efficient numerical computation. Computing in science & engineering, 13(2), 22\u201330. URL: https://doi.org/10.1109/MCSE.2011.37, doi:10.1109/MCSE.2011.37

Virtanen, P., Gommers, R., Oliphant, T. E., Haberland, M., Reddy, T., Cournapeau, D., \u2026 others. (2020). Scipy 1.0: fundamental algorithms for scientific computing in python. Nature methods, 17(3), 261\u2013272. URL: https://doi.org/10.1038/s41592-019-0686-2, doi:10.1038/s41592-019-0686-2
